# Insights into the metabolic consequences of type 2 diabetes

**DOI:** 10.1101/2024.06.20.599832

**Authors:** Ozvan Bocher, Archit Singh, Yue Huang, Urmo Võsa, Ene Reimann, Ana Arruda, Andrei Barysenska, Anastassia Kolde, Nigel W. Rayner, Estonian Biobank research team, Tõnu Esko, Reedik Mägi, Eleftheria Zeggini

## Abstract

Circulating metabolite levels have been associated with type 2 diabetes (T2D), but the extent to which these are affected by T2D and the involvement of genetics in mediating these relationships remain to be elucidated. In this study, we investigate the interplay between genetics, metabolomics and T2D risk in the UK Biobank dataset. We find 79 metabolites with a causal association to T2D, mostly spanning lipid-related classes, while twice as many metabolites are causally affected by T2D liability, including branched-chain amino acids. Secondly, using an interaction quantitative trait locus (QTL) analysis, we describe four metabolites, consistently replicated in an independent dataset from the Estonian Biobank, for which genetic loci in two different genomic regions show attenuated regulation in T2D cases compared to controls. The significant variants from the interaction QTL analysis are significant QTLs for the corresponding metabolites in the general population, but are not associated with T2D risk, pointing towards consequences of T2D on the genetic regulation of metabolite levels. Finally, we find 165 metabolites associated with microvascular, macrovascular, or both types of T2D complications, with only a few discriminating between complication classes. Of the 165 metabolites, 40 are not causally linked to T2D in either direction, suggesting biological mechanisms specific to the occurrence of complications. Overall, this work provides a map of the metabolic consequences of T2D and of the genetic regulation of metabolite levels and enable to better understand the trajectory of T2D leading to complications.

## Introduction

Type 2 diabetes (T2D) is a common, complex disease, with a prevalence that is expected to increase dramatically, and for which the genetics has been largely described through genome-wide association study (GWAS) meta-analysis efforts^1–3^. The next step towards translating these associations into the clinic is to understand the biological mechanisms behind them. To this end, multi-omics data such as metabolite levels offer great promise, as they enable the study of molecular phenotypes closely implicated in the disease. A large number of metabolites have been associated with T2D, as highlighted in a recent study from Julkunen et al.^4^, which described 230 metabolites as nominally associated with incident and prevalent T2D. Consistent associations between T2D risk and metabolites across studies mostly cover increases in various amino acid levels, especially branched-chain amino acids (BCAAs)^5–7^, and dyslipidemia^4, 8^. However, the predictive value of metabolite profiles in T2D risk is still debated. Improved prediction of various complex traits using metabolite profiles over genetic scores has been shown^9^, but this improvement is rather limited when compared to classical risk factors including age, sex and family history of disease^10, 11^. The limited prediction of metabolite profiles may be related to the uncertain causal role in modulating T2D risk, a question investigated using Mendelian randomization (MR)^12^. One particular example is BCAAs, for which some studies have described a causal effect of valine, leucine and/or isoleucine on T2D^13–15^, while more recent evidence has shown no causal effect of BCAAs on T2D risk in the UK Biobank (UKBB) cohort^16^. Conversely, increasing evidence of a causal effect of T2D liability on metabolite profiles has been shown, with increased alanine levels caused by increased T2D liability being consistently reported^13, 17, 18^. Conflicting conclusions exist for other metabolites, and little is known about the role of genetics in mediating these relationships. The genetic regulation of metabolite profiles is increasingly described, for example through the latest meta-analysis reporting over 400 independent loci associated with 233 metabolite levels^19^. These metabolite QTLs, also called mQTLs, are usually reported in the general population, and studies are starting to emerge on how they are modified by factors such as diet^20^. However, no study to date has investigated the question of whether the genetic regulation of metabolite levels is affected by the occurrence of T2D.

A large part of the healthcare burden associated with T2D is due to subsequent complications of the disease. T2D complications span both microvascular complications, which refer to complications involving small vessels and have an estimated prevalence of 53%, and macrovascular complications which refer to complications involving large vessels such as arteries and veins with an estimated prevalence of 27%^21^. Increasing evidence is emerging that metabolite profiles are also associated with the risk of developing complications, such as the fatty acid biosynthesis pathway with retinal and renal complications^22^, two of the main microvascular complications of T2D, n-3 fatty acids with T2D macrovascular complications^23^, or amino acids with both types of T2D complications^24^. However, replication of these associations is still needed, as well as investigating whether the metabolites associated with T2D complications are distinct from the ones causally affected by T2D liability. Exploring the links between metabolite levels and T2D can help us gain insights into how these relationships may influence trajectories towards T2D complications, for which underlying biological mechanisms are still to be unraveled.

Here, we aim to address these questions and elucidate the metabolic consequences of T2D by investigating the links between metabolite profiles, genetics, and the risk of T2D and subsequent complications. For this, we considered profiles of 249 metabolites, characterized for almost 275,000 participants from the UKBB cohort^25^ through two releases, which were here considered for discovery (n=117,967 individuals) and replication (n=156,385 individuals), or meta-analyzed when statistically appropriate. We first performed a bi-directional two sample MR analysis, using non-overlapping datasets for the SNP-exposure and SNP-outcome associations to limit potential bias^12^. Previous MR studies performed in the UKBB cohort have used only the first release of data for the 249 metabolites and were either limited to a few metabolites, or only investigated a single causal direction^16, 18, 26^. Secondly, we performed an interaction mQTL analysis to assess whether there is a different genetic regulation of metabolite levels between T2D patients and controls. Finally, we investigated how metabolite profiles are associated with T2D complications, and whether these associated metabolites overlap with the metabolites causally affected by T2D liability.

## Methods

### Data and quality control

Genetic data from genotyping arrays and imputation are available for over 500,000 participants in the UKBB cohort. Quality control (QC) was performed at the variant level and at the sample (S1 File). We selected only variants with a minor allele frequency greater than 1% and with an imputation INFO score greater than 0.8. To maximize sample homogeneity and to minimize potential bias in MR due to differences between the exposure and outcome datasets, we chose to focus on individuals of European ancestry. In total, 408,194 individuals and 9,572,578 variants remained after QC.

A total of 249 metabolite levels obtained from the Nightingale platform using nuclear magnetic resonance (NMR) spectroscopy is available in the UKBB cohort as part of two different releases covering a total of 274,352 individuals: 117,967 in the first release, and 156,385 in the second release. Absolute concentrations cover 168 metabolites, with an additional set of 81 metabolite ratios and percentages, which are further derived from the absolute concentrations (https://biobank.ndph.ox.ac.uk/showcase/refer.cgi?id=1682). To perform metabolite QC, we used the ukbnmr R package (version 1.5) specifically developed to remove technical variations from NMR metabolite measurements in the UKBB^27^. For each release, we considered only metabolite data at the first timepoint, T0, corresponding to a total of 227,607 individuals and 249 metabolites after applying the ukbnmr QC (S1 File). All analyses have been performed on inverse normal transformed values to obtain normally distributed metabolite levels^28^, which correspond to ‘metabolite levels’ for the rest of the manuscript.

### Definition of phenotypes

#### T2D status

T2D status was defined based on the UKBB field 130708, which corresponds to the first date of T2D report (self-reported or ICD10 code). Status was defined at T0 to analyze metabolite profiles in the light of T2D status at the time of profiling (S1 Fig), prevalent cases corresponding to individuals having a T2D diagnosis date before the date of blood sample collection on which metabolites were assayed, and incident cases to individuals having a T2D diagnosis date after the date of blood sample collection. Considering that T2D is often diagnosed with a few years delay^29^, we considered HbA1c levels in addition of the field 130708 to recover individuals likely having undiagnosed T2D at T0 among T2D incident cases. We removed from the analysis individuals with any mention of T1D (ICD10 code E10*, field 130706) or gestational diabetes (ICD10 code O24*, field 132202), and individuals diagnosed with T2D before the age of 36 years, in accordance with previous guidelines^30^. Individuals with a mention of T1D or gestational diabetes were also removed from the controls. The final number of T2D cases and controls included in the analyses, passing the genetic data QC and having measured metabolite levels are 3,088 and 88,244 for the first release and 4,302 and 118,555 for the second release, respectively.

#### T2D complications

We categorized prevalent cases of T2D at T0 into four mutually exclusive complication groups based on the occurrence of generalized vascular complication events: “microvascular”, “macrovascular”, “micro and macrovascular” complications, and “no complications”. “Microvascular” and “macrovascular” complications were defined based on ICD10 codes summarized in S1 Table. Individuals with an ICD10 code for both types of complications were attributed to the “micro and macrovascular” group and removed from the two other complication groups. We selected individuals with a diagnosis date of complications before the date of metabolite sample collection but after the T2D diagnosis date. If individuals had multiple ICD10 codes for the “microvascular” or the “macrovascular” complication group, the date of the first event was considered. For the “micro and macrovascular” group, the latest date between the “microvascular” and the “macrovascular” onset was considered. The total number of individuals in each complication group assayed in each of the release datasets is presented in Table 1.

**Table 1:**
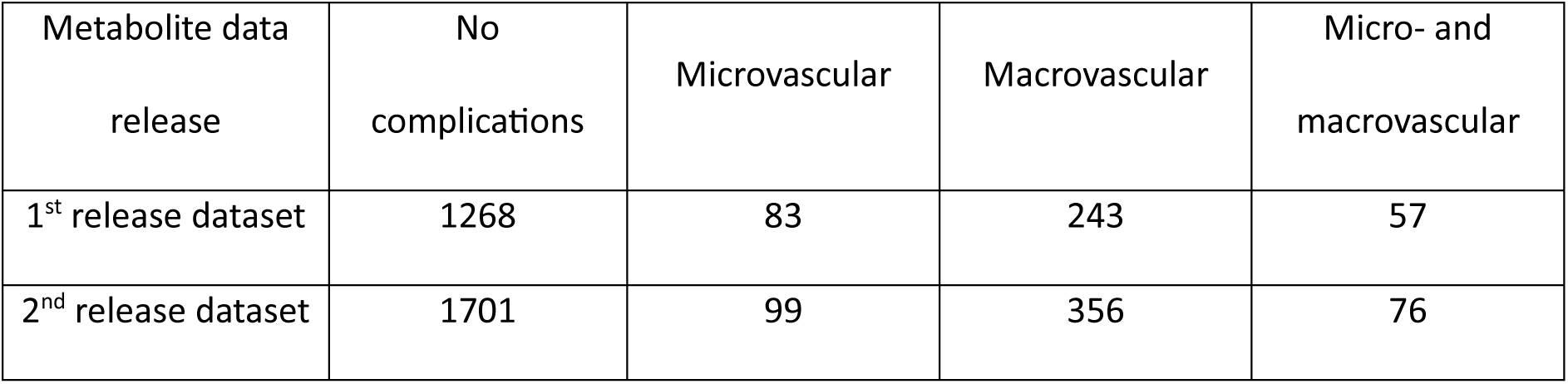
Number of T2D patients per complication group for each of the released dataset.

| Metabolite data<br>release | No<br>complications | Microvascular | Macrovascular | Micro- and<br>macrovascular |
| --- | --- | --- | --- | --- |
| 1<sup>st</sup> release dataset | 1268 | 83 | 243 | 57 |
| 2<sup>nd</sup> release dataset | 1701 | 99 | 356 | 76 |

### Statistical analyses

#### General considerations

All the analyses have been performed using R version 4.2.0 and adjusted for age, sex, and fasting time (only available in the UKBB), as well as for additional covariates in the analyses involving genetic data (S2 Table). To correct our analyses for multiple testing, we considered FDR-adjusted p-values (Benjamini-Hochberg method, referred to as q-values) to assess significance at 5%. All analyses were performed on the two release datasets separately and meta-analyzed, except for the MR analyses where the same T2D summary statistics were used for both MR analyses on the two release sets (details below).

#### Mendelian randomization

Two sample bi-directional MR was performed to assess the causal effects of metabolite levels on T2D risk (forward MR) and the causal effects of T2D liability on metabolite levels (reverse MR). MR was run separately on the two released datasets as it was not possible to perform meta-analysis due to the same outcome data being used in the two MR analyses, namely T2D DIAMANTE meta-analysis. For the first release dataset, we used mQTL summary statistics from Borges et al.^31^, and for the second release, we performed a mQTL analysis using the REGENIE software (version 2.2.4)^32^. For the associations between genetic variants and T2D, we used the DIAMANTE 2018 meta-analysis^2^, restricted to European ancestry samples, and without the UKBB cohort to avoid sample overlap between the exposure and the outcome data.

Instrumental variables (IVs) were selected by first defining independent variants through LD-based clumping using plink^33^ with the following parameters: R^2^<0.001 in windows of 10Mb, a p-value threshold of 2.54×10^-10^ for the metabolite levels and of 5×10^-8^ for T2D. The threshold of 2.54×10^-10^ is a genome-wide threshold corrected by the number of effective tests^34^, estimated at 197, which enables to correct the analyses for multiple testing while considering the correlation between the metabolites. The strength of each IV was determined using F-statistics. For the first released dataset and T2D summary statistics, the F-statistic was estimated using 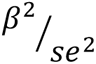, where beta is the effect estimate and se its corresponding standard error. For the second release dataset, individual-level data from the UKBB and the R package ivreg (version 0.6-1) were used to calculate the F-statistic. Only IVs with an F-statistic larger than 10 were retained.

MR was run using the R package TwoSampleMR^35^. RadialMR^36^ was used to assess heterogeneity and remove outliers for metabolites with a significant SNP heterogeneity. MR significance in the first release dataset was assessed based on the inverse variance weighted (IVW) method using q-values at a 5% threshold. Six additional MR methods (MR Egger, weighted median, simple mode, weighted mode, IVW fixed effect, IVW random effect and Steiger filtered IVW) were used as sensitivity analyses to check for concordant direction of causal effect estimate. We only considered metabolites having a non-significant heterogeneity and pleiotropy estimates, measured by the Q-statistic and the MR-Egger intercept, respectively. Significant metabolites were considered as replicated if they had an IVW q-value lower than 5% in the second release dataset with a concordant direction of effect of the IVW method with the first release dataset.

#### Interaction QTL

We performed an interaction QTL analysis to investigate the genetic regulation of metabolite levels in T2D cases and controls using the REGENIE software (version 2.2.4)^32^ and the following interaction test:

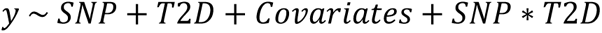

Where *SNP* ∗ *T2D* represents the interaction term between T2D disease status and genotypes. Only the p-value of the interaction test (“ADD-INT_SNPxVAR” column from the REGENIE output) was assessed for statistical significance. Interaction analyses were run on each of the two release datasets separately and were then meta-analyzed using the software METAL^37^ to maximize statistical power. We considered suggestive variants if they had a meta-analysis p-value lower than the genome-wide significance threshold of 5×10^-8^, a nominally significant p-value in both release datasets, and a concordant direction of effect between both datasets. To investigate whether the results from the interaction QTL analyses could be mediated by the effect of confounding variables, we performed sensitivity analyses where we sequentially adjusted for BMI, lipid-lowering medication, and metformin (File S1).

##### Replication of interaction QTL effects in Estonian Biobank

We replicated the interaction QTL analysis in the Estonian Biobank (EstBB), in which the same metabolite panel has been assayed, for the significant variants identified in the UKBB. The EstBB data freeze including altogether 211,728 biobank participants was applied. Individual level data analysis in the EBB was carried out under ethical approval [nr 1.1-12/3337] from the Estonian Committee on Bioethics and Human Research (Estonian Ministry of Social Affairs), using data according to release application [nr 6-7/GI/15486 T17] from the Estonian Biobank.

The T2D cases were defined using the same approach as in the UKBB, where T2D diagnosis was based on the ICD10 codes E11* from electronic health registries (EHRs), and further on HbA1c levels for incident T2D cases. Individuals who have consumed insulin at least one year after the T2D diagnosis were excluded (EHRs do not have information about prescription of Insulins and analogies (the Anatomical Therapeutic Chemical code A10A*)). The control group was defined as follows: (1) they do not have ICD10 codes E10*, E11* or O24* marked in their EHRs, (2) they have not been prescribed any of the drugs with the following ATC codes: A10A*, A10BA02, A10BF01, A10BB07, A10BB03, V04CA01, A10BB01, A10BB09, A10BB12, A10BG01, A10BX02, A10BG03, A10BX03, A10BG02.

Similar to UKBB, metabolite data is available from Nightingale NMR spectrometry platform, and a QC was performed on both these data and the genotyping data (S1 File). After QC, 6,237 T2D cases and 92,381 controls were enrolled into the replication analysis. Replication interaction QTL analysis was conducted by fitting a linear model with the interaction term between SNP and T2D status, while using SNP dosage, T2D status, sex, age at agreement, spectrometer serial number and ten first genetic principal components as covariates on each pair of variant and metabolite. Analysis and data processing was implemented into custom scripts and by using R v4.3.1. We considered replicated signals those with a q-value lower than 5% in the EstBB cohort and with a concordant direction of effect.

#### Analysis of T2D complications

We analyzed the differences between metabolite levels and the four complication groups defined previously (microvascular, macrovascular, micro and macrovascular, no complications) using a multinomial approach. We used the R package mlogit (version 1.1-1), with T2D individuals without complications representing the reference level. To increase statistical power, we meta-analyzed the results across the two release datasets for each complication group. We declared metabolites as having a significant effect between different complication groups if they had a q-value lower than 5% in the meta-analysis, and a nominal significant p-value in each of the release datasets with a concordant direction of effect. Finally, we performed forward one-sample MR analyses within the UKBB to assess the causal effect of metabolite levels on the risk of developing T2D complications using the R package ivreg (version 0.6-1). We used the same metabolite IVs as for the T2D bi-directional two-sample MR, and only kept the ones having an F-statistic greater than 10 in the subset of T2D individuals.

## Results

### Causal associations

Causal relationships between T2D and metabolites from the bi-directional MR that are significant in the first release dataset and replicated in the second release dataset are reported as an upset plot in Fig 1. This plot also includes the significant associations between metabolite levels and prevalent/incident T2D in the UKBB described by Julkunen et al.^4^ using the first metabolite release dataset. All metabolites that show a significant causal association in either MR direction in our analysis are also associated with prevalent or incident T2D reported by Julkunen et al.^4^, supporting the observed MR effects. Interestingly, while among the metabolites significant in the reverse direction, i.e., causally affected by T2D liability, 172 are found to be associated with both incident and prevalent T2D, 8 are found to be associated only with incident T2D and not significant in the forward direction. This finding points to T2D predisposition affecting these 8 metabolites levels, which are all related to very low-density lipoproteins (VLDL). MR estimates from the IVW method in both directions are reported in S3 Table and shown in Fig 2 as circular plots for 168 absolute metabolite levels and 81 derived ratios.

**Figure 1:**
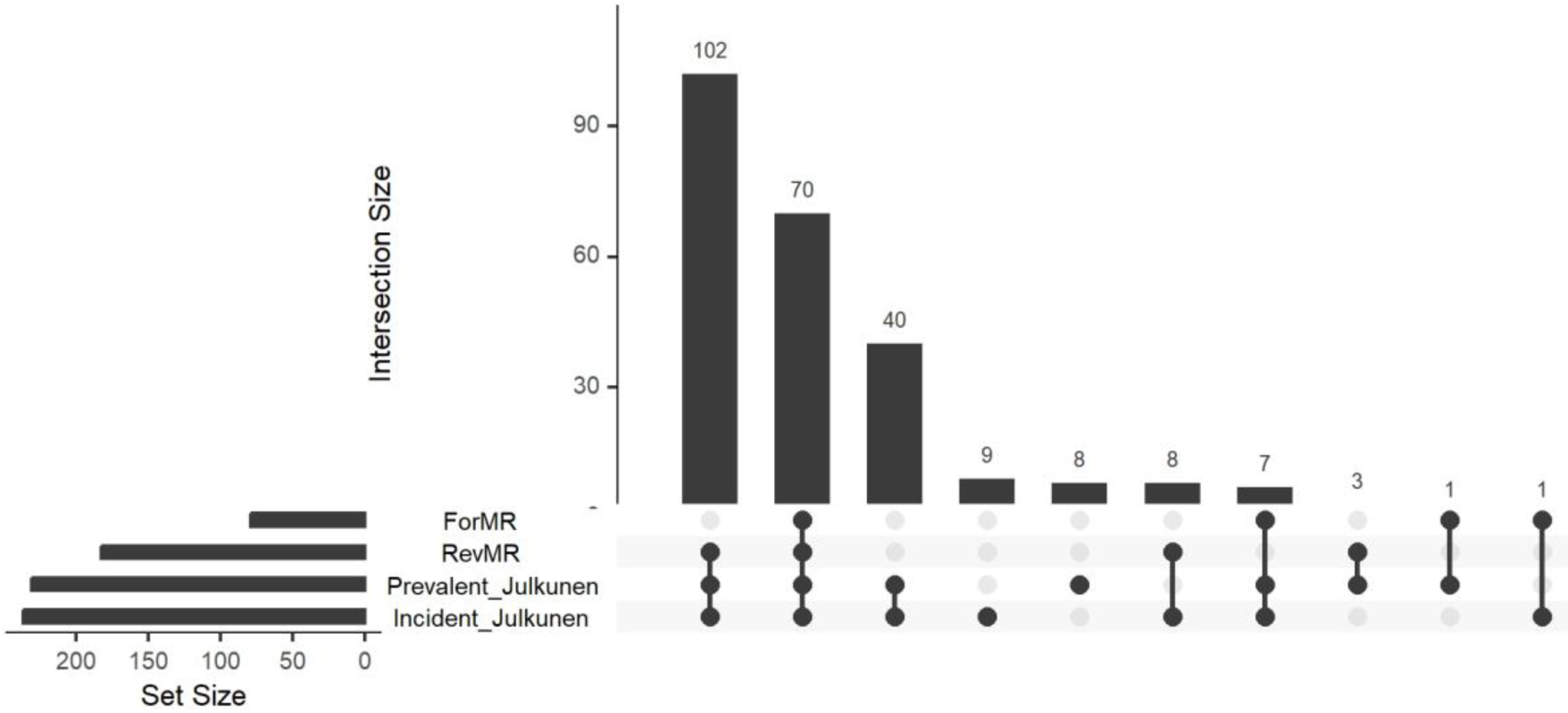
Upset plot of the forward and reverse MR analyses, along with the association results from Julkunen et al. between metabolite levels and prevalent/incident T2D.

**Figure 2:**
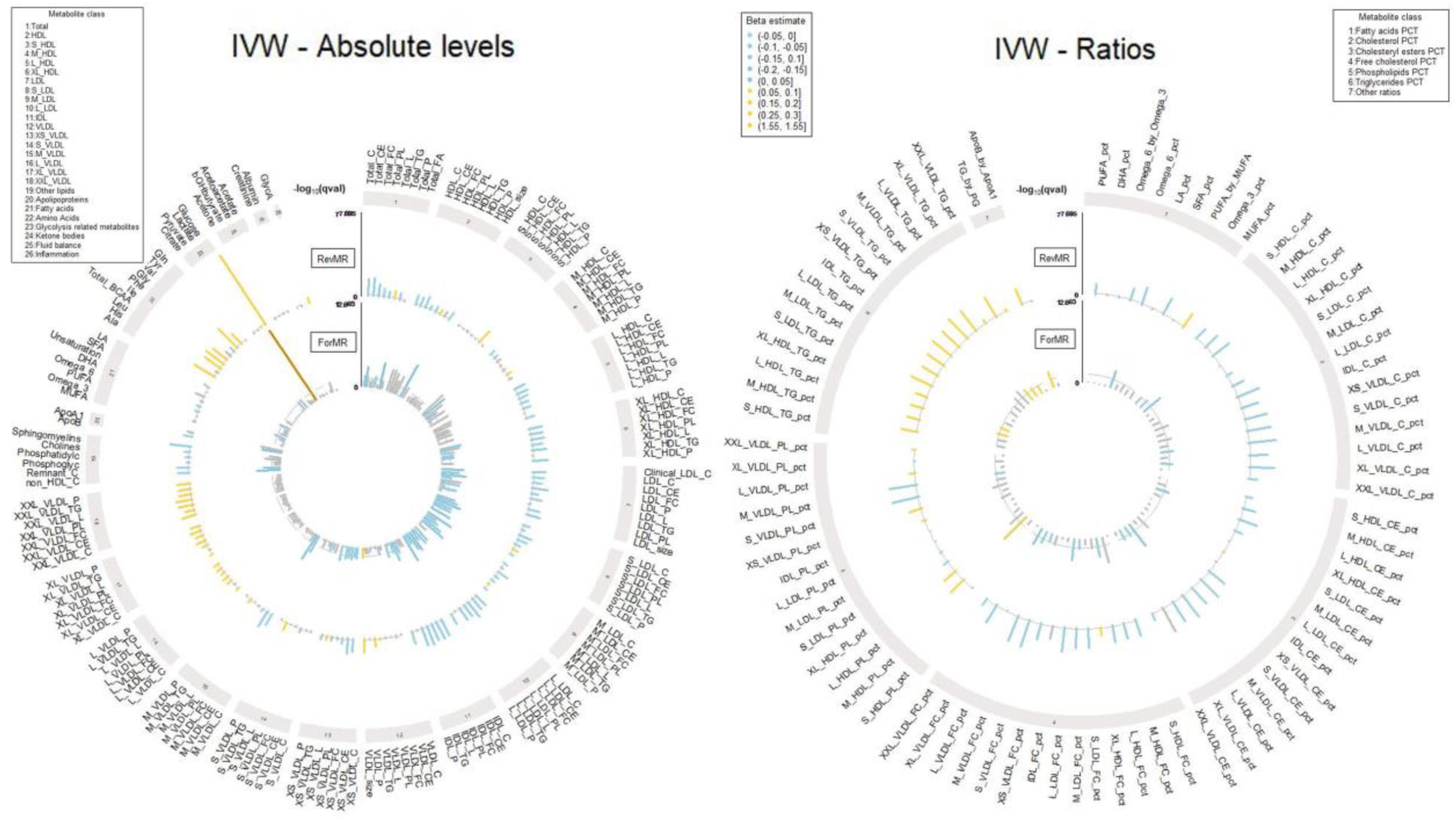
Circular plot of the forward and reverse MR analyses with estimates from the 1^st^ metabolite set. Metabolites having increased levels associated with T2D occurrence are presented in yellow and the ones with decreased levels in blue. Non-significant metabolites, or metabolites not replicated in the 2^nd^ metabolite set are colored in grey. Metabolites are grouped according to metabolite classes.

We identify 78 metabolites that are significant in the forward direction, i.e., that show a causal association with T2D risk, mostly spanning lipid classes, especially low-density lipoproteins (LDL) with decreased levels increasing the risk of T2D. While counterintuitive, increasing evidence suggests that individuals with low LDL-C have indeed a higher risk of developing T2D^38–40^, and a recent study described genetic variants associated with both higher T2D risk and lower LDL levels^41^. The LDL and T2D associations described here are also concordant with previous work using data from the UKBB^4, 18^. Glucose was the strongest signal in the first dataset (OR = 4.57 [3.15-6.63], p-value=8.73×10^-16^), which was replicated in the second released dataset. Additionally, some ratios in high density lipoproteins (HDL) and VLDL classes are found to be causally associated with T2D risk. We find no evidence of a causal role of absolute triglycerides (TG) levels on T2D, although it has been described as a predictor of T2D risk^17, 42^. We replicate findings from Mosley et al.^16^, which do not find evidence of a causal role of BCAAs on the risk of T2D. Almost all of the significant metabolites in the forward direction are also found to be associated in the reverse direction, highlighting the complex interplay between metabolite profiles and T2D liability.

We find 183 metabolites to be significantly associated in the reverse direction, of which 114 are not found in the forward direction. This includes a negative effect of T2D liability on large and very large HDL, intermediate-density lipoproteins (IDL) and LDL, while the opposite trend is observed for large and extremely large VLDL. Consistent trends are observed for lipoprotein ratios with T2D liability being causally associated with decreases in cholesterol and cholesteryl esters fractions, and with increases in TG ratios. Fatty acids percentages were also affected by T2D liability, with decreases in PUFA and omega-6 percentages, but increases in MUFA percentage. Finally, we describe causal effects of T2D liability on all BCAAs, as well as on tyrosine and alanine. We further compared our results with a reverse MR study that was carried out by Smith et al. on the first release dataset of metabolites from the UKBB^18^ using T2D summary statistics that included UKBB samples, and observe a high correlation of the effect estimates and p-values with our study (S2 Fig). Here, we report replication of 93% of the findings from Smith et al. in the second release dataset of metabolites from the UKBB without a potential bias due to sample overlap (S3 Table).

### Interaction QTL

We investigated whether causal effects of T2D liability on metabolite profiles could be mediated by a different genetic regulation of metabolite levels between T2D cases and controls through an interaction mQTL analysis. We identify 14 metabolites with variant having a significant interaction with T2D status, including glycine and various lipids. Of these, 40 variants were replicated in the EstBB (S4 Table), corresponding to four metabolites: percentage of free cholesterol in small HDL (S_HDL_FC_pct), percentage of phospholipids in large LDL (L_LDL_PL_pct), percentage of triglycerides in large VLDL (L_VLDL_TG_pct) and percentage of free cholesterol in large VLDL (L_VLDL_FC_pct). These significant interactions map to two intergenic regions, one shared by all metabolites except S_HDL_FC_pct, and the second showing significant genetic interactions with T2D status only for S_HDL_FC_pct (S3 Fig). The genetic associations of these variants with metabolite levels are stronger in the control group than in the T2D patients’ group, with some of the variants in the interaction regions being a significant mQTL only in the control group. For instance, rs6073958 is associated with lower levels of S_HDL_FC_pct in the controls group (beta = −0.146, p<1e-300), but not in the patients’ group (beta = −0.002, p = 0.93) (Fig 3).

**Figure 3:**
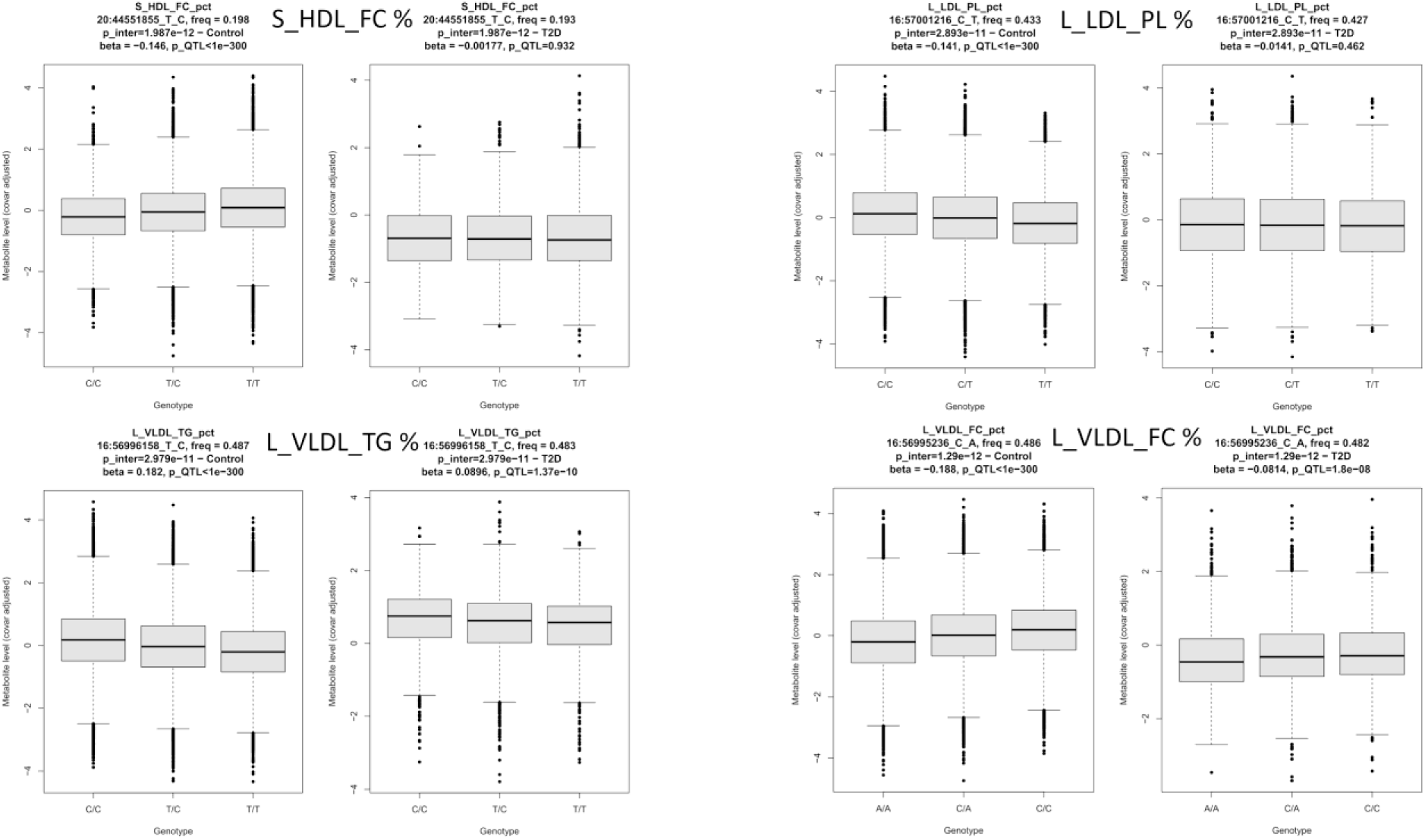
Boxplot showing metabolite levels according to the genotype in T2D patients and controls (the most significant variant for each of the four metabolites replicated in the EBB are represented). The metabolite levels represented are inverse-normal transformed and adjusted for the covariates used in the interaction QTL analyses. For each variant, its frequency in the control and in the T2D patients’ group is provided, along with the beta and p-value of association with metabolite levels. The overall p-value of the interaction test is also given.

L_VLDL_TG_pct is significant in both MR directions, while S_HDL_FC_pct and L_VLDL_FC_pct are only significant in the forward and reverse MR analysis respectively (Fig 4). The reverse MR estimates show evidence that T2D liability is causally associated with a decrease in S_HDL_FC_pct, and the major allele of the interacting variants are associated with increased levels of this metabolite. The opposite trend is observed for L_VLDL_TG_pct. These findings suggest that the causal effect of T2D liability identified in the reverse MR might be, at least partly, due to a genetic dysregulation of metabolite levels following T2D occurrence.

**Figure 4:**
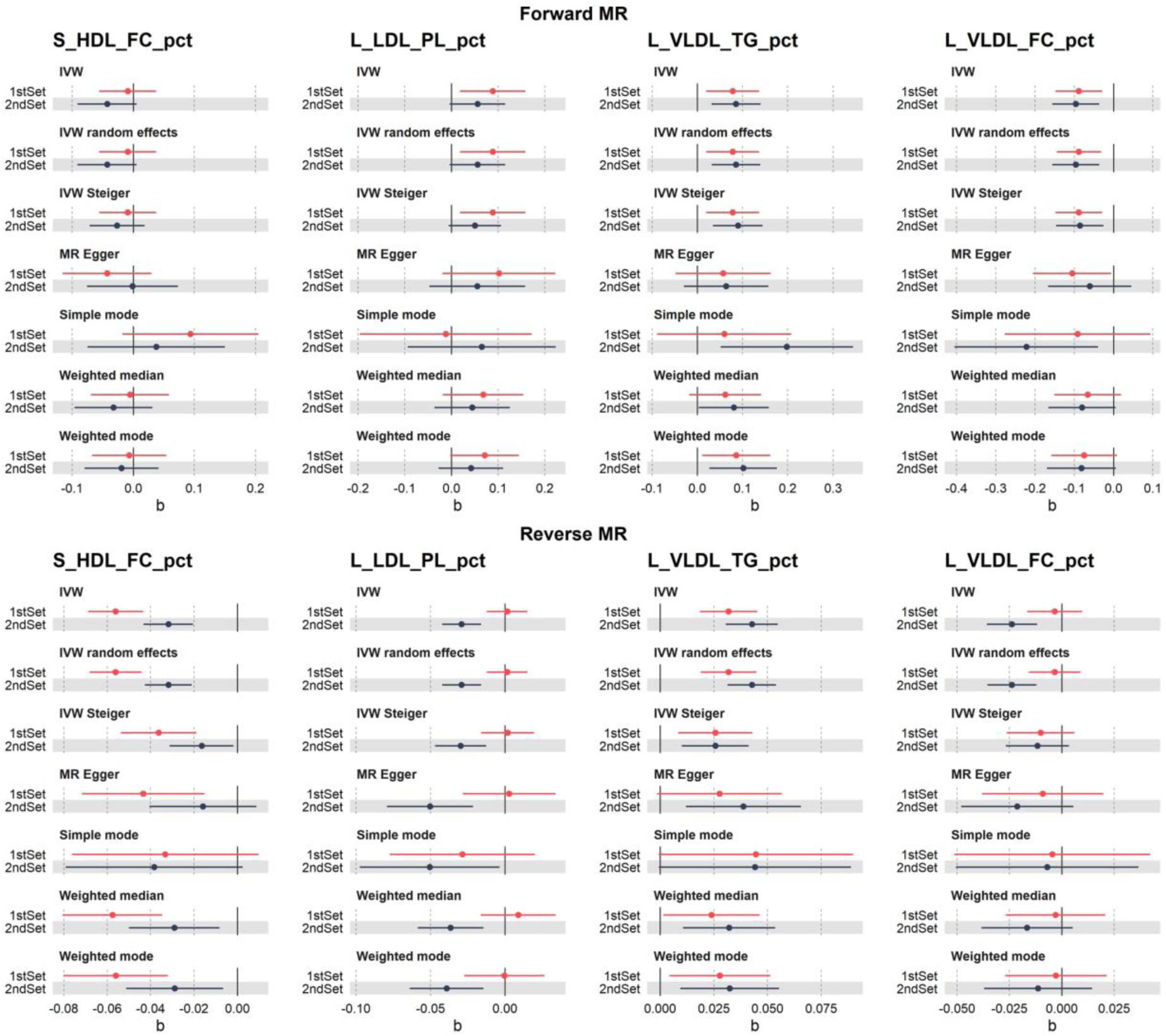
Bi-directional MR results for the four metabolites significant in the interaction QTL analysis. The top row corresponds to the forward MR (effects of metabolites on T2D), and the bottom row to the reverse MR (effects of T2D liability on metabolites). Effect sizes and standard error using IVW and the sensitivity methods are represented in red for the first set of metabolites, and in black for the second set of metabolites.

To gain more insights into the link between the four replicated metabolites and T2D, we looked at the most significant interaction QTLs. The most significant interaction QTL for L_VLDL_TG_pct, rs3816117, is annotated by the variant effect predictor (VEP^43^) as a modifier variant for cholesteryl ester transfer protein, (CETP, S4 Fig), a protein targeted by drugs to increase HDL-C and decrease LDL-C levels. The significant interaction variant for S_HDL_FC_pct, rs6073958, is annotated as a modified lncRNA. All of the interaction variants replicated in the EstBB are mQTLs for the corresponding metabolites in the overall UKBB cohort, but are not associated with T2D risk in the latest T2D GWAS meta-analysis^3^. These findings suggest that the different effect of genetics on the metabolite levels observed between T2D patients and controls is a consequence rather than a cause of the disease, in line with the results of the reverse MR for S_HDL_FC_pct and L_VLDL_TG_pct. The results remained consistent after adjustment for BMI, lipid medication or metformin medication (S5 Table).

### Metabolite profiles and T2D complications

We sought to better understand whether alterations in metabolite levels causally affected by T2D liability are associated with the risk of developing T2D complications (comparing T2D patients without complications, with microvascular complications, with macrovascular complications, and with both types of complications). Out of the 249 tested metabolites, 165 (66%) are associated with at least one of the complication groups, of which 156 are associated with the macrovascular group (Fig 5A). Only 3 and 6 signals are exclusive to the ‘microvascular’ and ‘both complications’ group, respectively. This pattern likely reflects the power of the analysis as the macrovascular group is three to five times larger than the two other complications groups (Table 1). Most of the significant metabolites are shared across multiple complication groups, showing that there is a metabolic signature associated with T2D complications.

**Figure 5:**
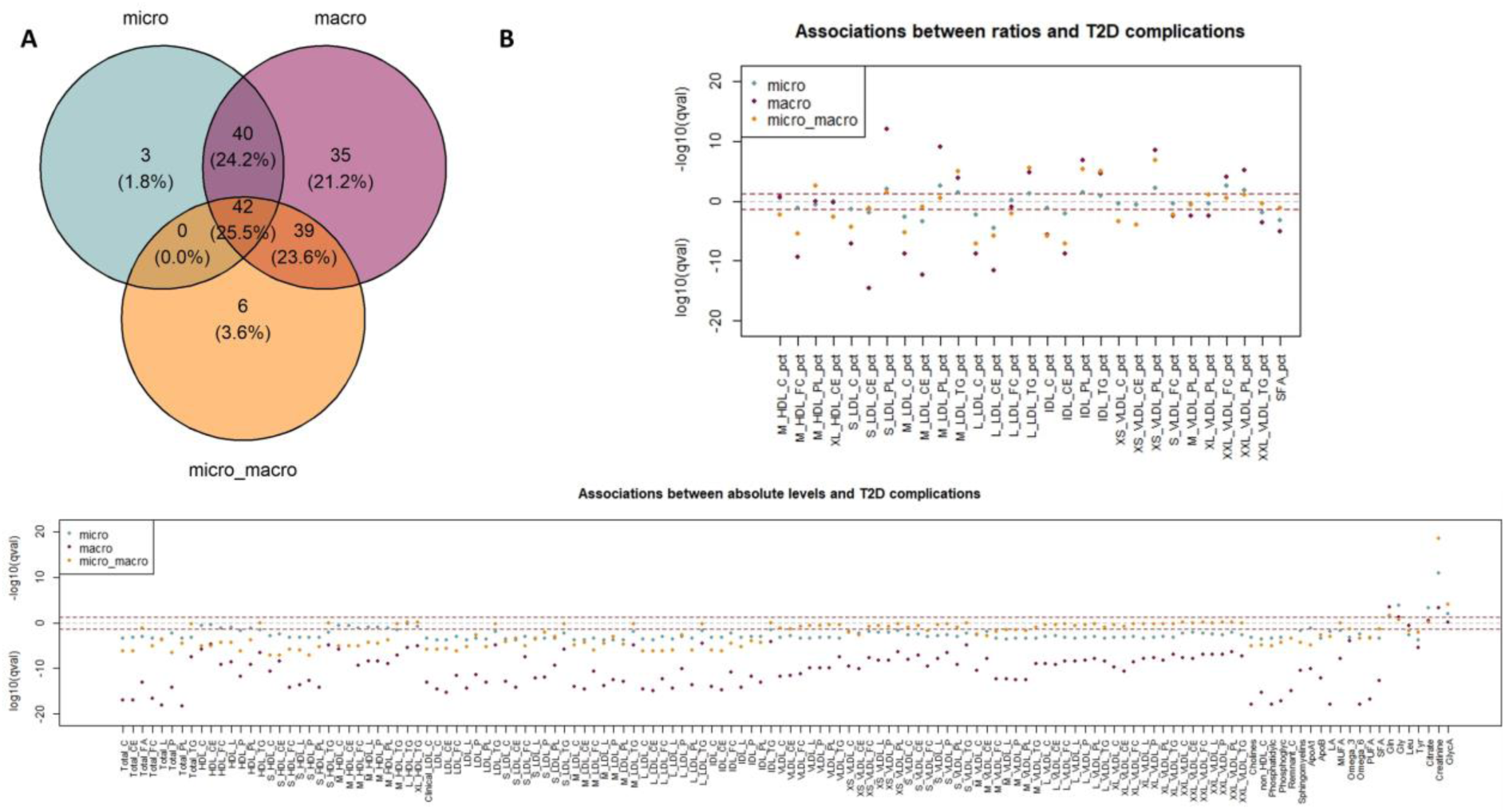
Association analyses between metabolite levels and T2D complications. A: Venn diagram of the significant metabolites (meta-analysis q-values lower than 5%) across complications. B: log10(q-values) of associations between metabolite levels (separated into absolute levels and derived ratios) and each complication group. –log10(q-values) are represented for positive associations, and log10(q-values) are represented for negative associations.

Four metabolites are more strongly associated with microvascular complications than with macrovascular complications. This is for example the case for creatinine, which is one of the strongest signals associated with microvascular complications (OR = 1.85 [1.58-2.18], p = 3.59×10^-14^). Creatinine is used in estimated glomerular filtration rate (eGFR) calculation, a measure used to assess kidney function, and known to be affected in nephropathy which is one of the main T2D microvascular complications^44^. Leucine is another example, for which significant association is found for the microvascular group (OR = 0.76 [0.65-0.89], p = 7.33×10^-4^), concordant with previous studies describing negative associations between leucine and kidney disease in T2D patients^45^. Increased levels of leucine were found to be putatively caused by an increased T2D liability in our reverse MR analysis, in line with the existing literature^6^, showing opposite associations of leucine with T2D and its complications. Overall, we find the profile of associations with metabolites to be similar with the MR estimates observed with T2D liability, with 78% of the metabolites associated with at least one complication and significant in the reverse MR having concordant direction of association between the two analyses, including decreases in most cholesterol metabolites, apolipoprotein B and glycine. Out of the 165 significantly associated metabolites, 54 and 118 are significant in the forward and reverse MR analyses, respectively. Of these, glycine and glycoprotein A are significant only in the reverse direction and associated with microvascular complications, suggesting that increased risk of these complications in T2D patients could be partly mediated by an impact of the T2D liability on metabolite levels. A total of 40 metabolites were associated with complications but not significant in the MR analyses, including creatinine, glutamine, and apolipoprotein A1, which were all previously associated with microvascular complications in the literature^44, 46, 47^. This finding points towards molecular mechanisms involving these metabolites being more specific to disease complications rather than T2D itself.

The metabolite associations with T2D complications stay stable upon adjustment, with 164, 158 and 156 signals remaining significant upon lipid medication, BMI and metformin adjustment, respectively (S6 Table). Metabolites affected by metformin and BMI adjustment span VLDL and HDL classes, while for instance valine, total VLDL size, albumin and total BCAAs become significant after adjustment. When applying one-sample MR between metabolite levels and the risk of T2D complications, we find no evidence of causal associations, potentially due to the low statistical power associated of the within-T2D UKBB MR analysis. Altogether, these results show that some metabolites causally affected by T2D liability are also associated with T2D complications, while others are specific to the development of complications, even though causality is still to be demonstrated.

## Discussion

In this study, we have investigated the links between plasma metabolite profiles, genetics, and the risk of T2D and subsequent complications, to evaluate the metabolic consequences of T2D. We have identified more metabolites as causally affected by T2D liability rather than having a causal effect on disease risk. Further, we have shown that the deregulation of metabolite levels following T2D occurrence may be partly due to different genetic effects in T2D patients and controls. Finally, we describe that metabolite profiles are also associated with T2D complications, though no significant causal relationship could be demonstrated in the present study.

Among the 183 metabolites causally affected by T2D liability, we found positive estimates with BCAAs, as well as with alanine and tyrosine, in agreement with previous studies^13, 18, 26^, but we did not find causal effects of BCAAs on T2D risk. Tyrosine was significant in both MR directions, but with opposite direction of effect, with decreased tyrosine levels being causally linked to increased T2D risk and T2D liability having a causal effect on increased tyrosine levels. These findings are in concordance with previous MR studies on the same cohort^26^ but not with observational studies, which describe positive associations between tyrosine levels and T2D risk^6^. Furthermore, we find negative associations between tyrosine levels and the three complications groups tested, in line with a previous study^24^, suggesting complex relationships between this metabolite and the risk of T2D complications. Further work is required to better disentangle the effect of tyrosine on T2D etiology. Glycine was significant only in the reverse MR analysis and associated with microvascular complications only. This amino acid has been already described as exhibiting lower levels in T2D patients, which could play a role in aggravating glucose dysregulation^48^. Even though not replicated in the EstBB, glycine presented the strongest signal in the interaction QTL analysis in the UKBB, with a weaker genetic regulation in T2D patients compared to controls. A genetic dysregulation of glycine levels might, therefore, be involved in T2D complications etiology, but replication is needed in an independent cohort.

Our findings also provide insights into the role of lipoprotein classes in T2D etiology. For example, TG levels have been described to be positively associated with T2D^41^ and are known to positively correlate with glucose levels, a relationship that may be exacerbated in individuals with high polygenic risk scores for T2D^49^. TG levels, which we found to be causally affected by T2D liability and for some of the related percentages significant in our interaction QTL analysis, may therefore contribute to increased glucose levels. For all other types of lipoproteins that were significant in the reverse MR analyses, T2D liability had a causal effect on decreasing their levels. This includes absolute LDL levels and to a lesser extent their related percentages, in line with previous studies^4, 18, 38–40^. However, these relationships are still debated, and caution is to be taken when interpreting results in studies not restricted to medication-free individuals. For example, Smith et al.^18^ showed in a similar reverse MR setting, using age as a proxy for medication use, that lipoprotein associations could be distorted upon medication use. Lipid-related medication adjusted T2D summary statistics are not available, and even if they were, care should be taken in interpreting results from such MR analyses using adjusted summary statistics due to potential collider bias^50^.

We identify 165 associations between metabolite levels and the risk of developing T2D complications, most of which are shared across complication groups with only few, such as creatinine and glycine, being more strongly associated with microvascular than macrovascular outcomes despite smaller sample size of the microvascular group. A total of 118 metabolites were also found to be significant in the MR analyses, with similar profiles of association. A counter example of this is leucine, which showed associations in opposite directions between the reverse MR analysis and the differential level analysis with T2D complications. The discrepancies observed in our study between the two MR directions, as well as with the risk of complications, highlight the need for further work to better understand disease trajectories of T2D and its complications. A total of 40 metabolites were associated with at least one complication group while not being significant in any of the MR analyses with T2D, suggesting mechanisms specific to T2D complications. However, causal inference to disentangle causation from association in the risk of T2D complications is warranted but challenging given the limited statistical power of MR analysis restricted to T2D patients.

In addition to investigating relationships between metabolite levels and T2D, we describe for the first time, to our knowledge, differences in their genetic regulation between T2D patients and controls for 14 metabolites, of which four were replicated in an independent cohort. All of the significant variants identified are found to be significant mQTLs in the overall UKBB but are not associated with T2D risk^3^. These results suggest that deregulated levels of these metabolites are a consequence rather than a cause of the disease, in line with the observations from the bi-directional MR, where evidence is found in the reverse direction for two of these four metabolites. We have performed an agnostic genome-wide scan to look for interaction signals. Given the results using this approach, we have observed that all the interaction variants, which are mQTLs at a population-level, lead to a decreased magnitude of metabolite genetic regulation in the T2D patients’ group compared to the control group. This could be of clinical relevance, as for example some of the interaction variants identified in our study are regulators of CETP, a target of lipid-lowering drugs, which have been shown to correlate with diabetes incidence^51^.

The present work presents some limitations. We have performed bi-directional MR using a similar framework to previous studies in the UKBB based on the first release dataset of metabolite data^16, 18^, with the benefit of providing replication in the second release dataset. The use of the same outcome data for the two release datasets prevented the meta-analysis of the results. In this study, we are restricted to the European ancestry DIAMANTE study from 2018 because it is the latest one with summary statistics available without the UKBB samples, which enables us to avoid sample overlap between the exposure and outcome and to limit potential related bias^12^. Our results need to be extended to non-European populations, which will require global efforts to characterize the genetic regulation of molecular traits in these populations, along with methodological developments to deal with multi-ancestry data, especially in MR studies. Additionally, our findings warrant replication in cohorts external to the UKBB, especially for the MR analyses and complication associations. Finally, our study provides useful insights into the metabolic consequences of T2D but is limited by the assayed metabolite panel, which is mostly composed of lipid-related plasma metabolites. Additional metabolite classes, as well as metabolite levels from different tissue types would help in better unravelling biological mechanisms.

While MR studies enable the assessment of causality between an exposure and an outcome, they are prone to false positives and based on the liability of the disease but not its occurrence. By using interaction models, we went one step further to describe the consequences of T2D occurrence on metabolite levels and their genetic regulation. We highlight that changes in metabolite profiles can be useful to better understand T2D disease progression, as exemplified by the metabolites associated with the risk of developing T2D complications. Altogether, our results enable a deeper understanding of the metabolic consequences of T2D and provide future directions for the study of the genetic regulation of molecular levels in T2D and its complications to better capture disease trajectory.

## Data availability

Summary statistics of the mQTL analysis in the second data release of the UKBB have been submitted to the GWAS catalog and will be released upon publication.

## Acknowledgements

This research has been conducted using the UK Biobank Resource under Application Number 10205.

Ozvan Bocher, Archit Singh, Yue Huang, Ene Reimann and Reedik Mägi have received funding from the European Union’s Horizon 2020 research and innovation programme under Grant Agreement No 101017802 (OPTOMICS).

Tõnu Esko and Urmo Võsa are supported by Estonian Research Council grant PUT (PRG1291) and Reedik Mägi by Estonian Research Council grant PUT (PRG1911) and by Estonian Ministry of Education and Research Funding (TK214).

The Estonian Biobank research team refers to Andres Metspalu, Lili Milani, Tõnu Esko, Reedik Mägi, Mari Nelis and Georgi Hudjashov.

Data analysis in the EstBB was carried out in part in the High-Performance Computing Center of University of Tartu. The activities of the EstBB are regulated by the Human Genes Research Act, which was adopted in 2000 specifically for the operations of the EstBB. Individual level data analysis in the EBB was carried out under ethical approval [nr 1.1-12/3337] from the Estonian Committee on Bioethics and Human Research (Estonian Ministry of Social Affairs), using data according to release application [nr 6-7/GI/15486 T17] from the Estonian Biobank.

## Notes

### Competing Interest Statement

The authors have declared no competing interest.

### Summary of Updates

Addition of the supplementary information (supplementary tables, supplementary figures, supplementary information).

## References

1. Mahajan, A., Spracklen, C.N., Zhang, W., Ng, M.C.Y., Petty, L.E., Kitajima, H., Yu, G.Z., Rueger, S., Speidel, L., Kim, Y.J., et al. (2022). Multi-ancestry genetic study of type 2 diabetes highlights the power of diverse populations for discovery and translation. Nat Genet 54, 560–572. 10.1038/s41588-022-01058-3.

2. Mahajan, A., Taliun, D., Thurner, M., Robertson, N.R., Torres, J.M., Rayner, N.W., Payne, A.J., Steinthorsdottir, V., Scott, R.A., Grarup, N., et al. (2018). Fine-mapping type 2 diabetes loci to single-variant resolution using high-density imputation and islet-specific epigenome maps. Nat Genet 50, 1505–1513. 10.1038/s41588-018-0241-6.

3. Suzuki, K., Hatzikotoulas, K., Southam, L., Taylor, H.J., Yin, X., Lorenz, K.M., Mandla, R., Huerta-Chagoya, A., Melloni, G.E.M., Kanoni, S., et al. (2024). Genetic drivers of heterogeneity in type 2 diabetes pathophysiology. Nature. 10.1038/s41586-024-07019-6.

4. Julkunen, H., Cichonska, A., Tiainen, M., Koskela, H., Nybo, K., Makela, V., Nokso-Koivisto, J., Kristiansson, K., Perola, M., Salomaa, V., et al. (2023). Atlas of plasma NMR biomarkers for health and disease in 118,461 individuals from the UK Biobank. Nat Commun 14, 604. 10.1038/s41467-023-36231-7.

5. Morze, J., Wittenbecher, C., Schwingshackl, L., Danielewicz, A., Rynkiewicz, A., Hu, F.B., and Guasch-Ferre, M. (2022). Metabolomics and Type 2 Diabetes Risk: An Updated Systematic Review and Meta-analysis of Prospective Cohort Studies. Diabetes Care 45, 1013–1024. 10.2337/dc21-1705.

6. Wang, T.J., Larson, M.G., Vasan, R.S., Cheng, S., Rhee, E.P., McCabe, E., Lewis, G.D., Fox, C.S., Jacques, P.F., Fernandez, C., et al. (2011). Metabolite profiles and the risk of developing diabetes. Nat Med 17, 448–453. 10.1038/nm.2307.

7. Sun, Y., Gao, H.Y., Fan, Z.Y., He, Y., and Yan, Y.X. (2020). Metabolomics Signatures in Type 2 Diabetes: A Systematic Review and Integrative Analysis. J Clin Endocrinol Metab 105. 10.1210/clinem/dgz240.

8. Jin, Q., and Ma, R.C.W. (2021). Metabolomics in Diabetes and Diabetic Complications: Insights from Epidemiological Studies. Cells 10. 10.3390/cells10112832.

9. Nightingale Health Biobank Collaborative, G., Jeffrey, C.B., Tõnu, E., Krista, F., Luke, J.-D., Pekka, J., Heli, J., Tuija, J., Nurlan, K., Sini, K., et al. (2023). Metabolomic and genomic prediction of common diseases in 477,706 participants in three national biobanks. medRxiv, 2023.2006.2009.23291213. 10.1101/2023.06.09.23291213.

10. Bragg, F., Trichia, E., Aguilar-Ramirez, D., Besevic, J., Lewington, S., and Emberson, J. (2022). Predictive value of circulating NMR metabolic biomarkers for type 2 diabetes risk in the UK Biobank study. BMC Med 20, 159. 10.1186/s12916-022-02354-9.

11. Buergel, T., Steinfeldt, J., Ruyoga, G., Pietzner, M., Bizzarri, D., Vojinovic, D., Upmeier Zu Belzen, J., Loock, L., Kittner, P., Christmann, L., et al. (2022). Metabolomic profiles predict individual multidisease outcomes. Nat Med 28, 2309–2320. 10.1038/s41591-022-01980-3.

12. Sanderson, E., Glymour, M.M., Holmes, M.V., Kang, H., Morrison, J., Munafo, M.R., Palmer, T., Schooling, C.M., Wallace, C., Zhao, Q., and Smith, G.D. (2022). Mendelian randomization. Nat Rev Methods Primers 2. 10.1038/s43586-021-00092-5.

13. Porcu, E., Gilardi, F., Darrous, L., Yengo, L., Bararpour, N., Gasser, M., Marques-Vidal, P., Froguel, P., Waeber, G., Thomas, A., and Kutalik, Z. (2021). Triangulating evidence from longitudinal and Mendelian randomization studies of metabolomic biomarkers for type 2 diabetes. Sci Rep 11, 6197. 10.1038/s41598-021-85684-7.

14. Lotta, L.A., Scott, R.A., Sharp, S.J., Burgess, S., Luan, J., Tillin, T., Schmidt, A.F., Imamura, F., Stewart, I.D., Perry, J.R., et al. (2016). Genetic Predisposition to an Impaired Metabolism of the Branched-Chain Amino Acids and Risk of Type 2 Diabetes: A Mendelian Randomisation Analysis. PLoS Med 13, e1002179. 10.1371/journal.pmed.1002179.

15. Yin, X., Li, J., Bose, D., Okamoto, J., Kwon, A., Jackson, A.U., Silva, L.F., Oravilahti, A., Stringham, H.M., Ripatti, S., et al. (2023). Metabolome-wide Mendelian randomization characterizes heterogeneous and shared causal effects of metabolites on human health. medRxiv. 10.1101/2023.06.26.23291721.

16. Mosley, J.D., Shi, M., Agamasu, D., Vaitinadin, N.S., Murthy, V.L., Shah, R.V., Bagheri, M., and Ferguson, J.F. (2024). Branched-chain amino acids and type 2 diabetes: a bidirectional Mendelian randomization analysis. Obesity (Silver Spring) 32, 423–435. 10.1002/oby.23951.

17. Liu, J., van Klinken, J.B., Semiz, S., van Dijk, K.W., Verhoeven, A., Hankemeier, T., Harms, A.C., Sijbrands, E., Sheehan, N.A., van Duijn, C.M., and Demirkan, A. (2017). A Mendelian Randomization Study of Metabolite Profiles, Fasting Glucose, and Type 2 Diabetes. Diabetes 66, 2915–2926. 10.2337/db17-0199.

18. Smith, M.L., Bull, C.J., Holmes, M.V., Davey Smith, G., Sanderson, E., Anderson, E.L., and Bell, J.A. (2023). Distinct metabolic features of genetic liability to type 2 diabetes and coronary artery disease: a reverse Mendelian randomization study. EBioMedicine 90, 104503. 10.1016/j.ebiom.2023.104503.

19. Karjalainen, M.K., Karthikeyan, S., Oliver-Williams, C., Sliz, E., Allara, E., Fung, W.T., Surendran, P., Zhang, W., Jousilahti, P., Kristiansson, K., et al. (2024). Genome-wide characterization of circulating metabolic biomarkers. Nature. 10.1038/s41586-024-07148-y.

20. Bar, N., Korem, T., Weissbrod, O., Zeevi, D., Rothschild, D., Leviatan, S., Kosower, N., Lotan-Pompan, M., Weinberger, A., Le Roy, C.I., et al. (2020). A reference map of potential determinants for the human serum metabolome. Nature 588, 135–140. 10.1038/s41586-020-2896-2.

21. Litwak, L., Goh, S.Y., Hussein, Z., Malek, R., Prusty, V., and Khamseh, M.E. (2013). Prevalence of diabetes complications in people with type 2 diabetes mellitus and its association with baseline characteristics in the multinational A1chieve study. Diabetol Metab Syndr 5, 57. 10.1186/1758-5996-5-57.

22. Tomofuji, Y., Suzuki, K., Kishikawa, T., Shojima, N., Hosoe, J., Inagaki, K., Matsubayashi, S., Ishihara, H., Watada, H., Ishigaki, Y., et al. (2023). Identification of serum metabolome signatures associated with retinal and renal complications of type 2 diabetes. Commun Med (Lond) 3, 5. 10.1038/s43856-022-00231-3.

23. Harris, K., Oshima, M., Sattar, N., Wurtz, P., Jun, M., Welsh, P., Hamet, P., Harrap, S., Poulter, N., Chalmers, J., and Woodward, M. (2020). Plasma fatty acids and the risk of vascular disease and mortality outcomes in individuals with type 2 diabetes: results from the ADVANCE study. Diabetologia 63, 1637–1647. 10.1007/s00125-020-05162-z.

24. Welsh, P., Rankin, N., Li, Q., Mark, P.B., Wurtz, P., Ala-Korpela, M., Marre, M., Poulter, N., Hamet, P., Chalmers, J., et al. (2018). Circulating amino acids and the risk of macrovascular, microvascular and mortality outcomes in individuals with type 2 diabetes: results from the ADVANCE trial. Diabetologia 61, 1581–1591. 10.1007/s00125-018-4619-x.

25. Sudlow, C., Gallacher, J., Allen, N., Beral, V., Burton, P., Danesh, J., Downey, P., Elliott, P., Green, J., Landray, M., et al. (2015). UK biobank: an open access resource for identifying the causes of a wide range of complex diseases of middle and old age. PLoS Med 12, e1001779. 10.1371/journal.pmed.1001779.

26. Jacky Man Yuen, M., Baoting, H., Tommy Hon Ting, W., Ying, L., Shan, L., Kenneth, L., Jimmy Chun Yu, L., and Shiu Lun Au, Y. (2023). Evaluating the role of amino acids in type 2 diabetes risk: a Mendelian randomization study. medRxiv, 2023.2008.2027.23294702. 10.1101/2023.08.27.23294702.

27. Ritchie, S.C., Surendran, P., Karthikeyan, S., Lambert, S.A., Bolton, T., Pennells, L., Danesh, J., Di Angelantonio, E., Butterworth, A.S., and Inouye, M. (2023). Quality control and removal of technical variation of NMR metabolic biomarker data in ∼120,000 UK Biobank participants. Sci Data 10, 64. 10.1038/s41597-023-01949-y.

28. Auer, P.L., Reiner, A.P., and Leal, S.M. (2016). The effect of phenotypic outliers and non-normality on rare-variant association testing. Eur J Hum Genet 24, 1188–1194. 10.1038/ejhg.2015.270.

29. Gopalan, A., Mishra, P., Alexeeff, S.E., Blatchins, M.A., Kim, E., Man, A.H., and Grant, R.W. (2018). Prevalence and predictors of delayed clinical diagnosis of Type 2 diabetes: a longitudinal cohort study. Diabet Med 35, 1655–1662. 10.1111/dme.13808.

30. Eastwood, S.V., Mathur, R., Atkinson, M., Brophy, S., Sudlow, C., Flaig, R., de Lusignan, S., Allen, N., and Chaturvedi, N. (2016). Algorithms for the Capture and Adjudication of Prevalent and Incident Diabetes in UK Biobank. PLoS One 11, e0162388. 10.1371/journal.pone.0162388.

31. Borges, M.C., Schmidt, A.F., Jefferis, B., Wannamethee, S.G., Lawlor, D.A., Kivimaki, M., Kumari, M., Gaunt, T.R., Ben-Shlomo, Y., Tillin, T., et al. (2020). Circulating Fatty Acids and Risk of Coronary Heart Disease and Stroke: Individual Participant Data Meta-Analysis in Up to 16 126 Participants. J Am Heart Assoc 9, e013131. 10.1161/JAHA.119.013131.

32. Mbatchou, J., Barnard, L., Backman, J., Marcketta, A., Kosmicki, J.A., Ziyatdinov, A., Benner, C., O’Dushlaine, C., Barber, M., Boutkov, B., et al. (2021). Computationally efficient whole-genome regression for quantitative and binary traits. Nat Genet 53, 1097–1103. 10.1038/s41588-021-00870-7.

33. Purcell, S., Neale, B., Todd-Brown, K., Thomas, L., Ferreira, M.A., Bender, D., Maller, J., Sklar, P., de Bakker, P.I., Daly, M.J., and Sham, P.C. (2007). PLINK: a tool set for whole-genome association and population-based linkage analyses. Am J Hum Genet 81, 559–575. 10.1086/519795.

34. Li, J., and Ji, L. (2005). Adjusting multiple testing in multilocus analyses using the eigenvalues of a correlation matrix. Heredity (Edinb) 95, 221–227. 10.1038/sj.hdy.6800717.

35. Hemani, G., Zheng, J., Elsworth, B., Wade, K.H., Haberland, V., Baird, D., Laurin, C., Burgess, S., Bowden, J., Langdon, R., et al. (2018). The MR-Base platform supports systematic causal inference across the human phenome. Elife 7. 10.7554/eLife.34408.

36. Bowden, J., Spiller, W., Del Greco, M.F., Sheehan, N., Thompson, J., Minelli, C., and Davey Smith, G. (2018). Improving the visualization, interpretation and analysis of two-sample summary data Mendelian randomization via the Radial plot and Radial regression. Int J Epidemiol 47, 1264–1278. 10.1093/ije/dyy101.

37. Willer, C.J., Li, Y., and Abecasis, G.R. (2010). METAL: fast and efficient meta-analysis of genomewide association scans. Bioinformatics 26, 2190–2191. 10.1093/bioinformatics/btq340.

38. Andersson, C., Lyass, A., Larson, M.G., Robins, S.J., and Vasan, R.S. (2015). Low-density-lipoprotein cholesterol concentrations and risk of incident diabetes: epidemiological and genetic insights from the Framingham Heart Study. Diabetologia 58, 2774–2780. 10.1007/s00125-015-3762-x.

39. Feng, Q., Wei, W.Q., Chung, C.P., Levinson, R.T., Sundermann, A.C., Mosley, J.D., Bastarache, L., Ferguson, J.F., Cox, N.J., Roden, D.M., et al. (2018). Relationship between very low low-density lipoprotein cholesterol concentrations not due to statin therapy and risk of type 2 diabetes: A US-based cross-sectional observational study using electronic health records. PLoS Med 15, e1002642. 10.1371/journal.pmed.1002642.

40. Besseling, J., Kastelein, J.J., Defesche, J.C., Hutten, B.A., and Hovingh, G.K. (2015). Association between familial hypercholesterolemia and prevalence of type 2 diabetes mellitus. JAMA 313, 1029–1036. 10.1001/jama.2015.1206.

41. Klimentidis, Y.C., Arora, A., Newell, M., Zhou, J., Ordovas, J.M., Renquist, B.J., and Wood, A.C. (2020). Phenotypic and Genetic Characterization of Lower LDL Cholesterol and Increased Type 2 Diabetes Risk in the UK Biobank. Diabetes 69, 2194–2205. 10.2337/db19-1134.

42. Zhao, J., Zhang, Y., Wei, F., Song, J., Cao, Z., Chen, C., Zhang, K., Feng, S., Wang, Y., and Li, W.D. (2019). Triglyceride is an independent predictor of type 2 diabetes among middle-aged and older adults: a prospective study with 8-year follow-ups in two cohorts. J Transl Med 17, 403. 10.1186/s12967-019-02156-3.

43. McLaren, W., Gil, L., Hunt, S.E., Riat, H.S., Ritchie, G.R., Thormann, A., Flicek, P., and Cunningham, F. (2016). The Ensembl Variant Effect Predictor. Genome Biol 17, 122. 10.1186/s13059-016-0974-4.

44. Levey, A.S., and Coresh, J. (2012). Chronic kidney disease. Lancet 379, 165–180. 10.1016/S0140-6736(11)60178-5.

45. Tofte, N., Vogelzangs, N., Mook-Kanamori, D., Brahimaj, A., Nano, J., Ahmadizar, F., van Dijk, K.W., Frimodt-Moller, M., Arts, I., Beulens, J.W.J., et al. (2020). Plasma Metabolomics Identifies Markers of Impaired Renal Function: A Meta-analysis of 3089 Persons with Type 2 Diabetes. J Clin Endocrinol Metab 105. 10.1210/clinem/dgaa173.

46. Rhee, S.Y., Jung, E.S., Park, H.M., Jeong, S.J., Kim, K., Chon, S., Yu, S.Y., Woo, J.T., and Lee, C.H. (2018). Plasma glutamine and glutamic acid are potential biomarkers for predicting diabetic retinopathy. Metabolomics 14, 89. 10.1007/s11306-018-1383-3.

47. Moosaie, F., Davatgari, R.M., Firouzabadi, F.D., Esteghamati, S., Deravi, N., Meysamie, A., Khaloo, P., Nakhjavani, M., and Esteghamati, A. (2020). Lipoprotein(a) and Apolipoproteins as Predictors for Diabetic Retinopathy and Its Severity in Adults With Type 2 Diabetes: A Case-Cohort Study. Can J Diabetes 44, 414–421. 10.1016/j.jcjd.2020.01.007.

48. Yan-Do, R., and MacDonald, P.E. (2017). Impaired “Glycine”-mia in Type 2 Diabetes and Potential Mechanisms Contributing to Glucose Homeostasis. Endocrinology 158, 1064–1073. 10.1210/en.2017-00148.

49. Lim, J.E., Kang, J.O., Ha, T.W., Jung, H.U., Kim, D.J., Baek, E.J., Kim, H.K., Chung, J.Y., Rhee, S.Y., Kim, M.K., et al. (2022). Gene-environment interaction in type 2 diabetes in Korean cohorts: Interaction of a type 2 diabetes polygenic risk score with triglyceride and cholesterol on fasting glucose levels. Genet Epidemiol 46, 285–302. 10.1002/gepi.22454.

50. Hartwig, F.P., Tilling, K., Davey Smith, G., Lawlor, D.A., and Borges, M.C. (2021). Bias in two-sample Mendelian randomization when using heritable covariable-adjusted summary associations. Int J Epidemiol 50, 1639–1650. 10.1093/ije/dyaa266.

51. Masson, W., Lobo, M., Siniawski, D., Huerin, M., Molinero, G., Valero, R., and Nogueira, J.P. (2018). Therapy with cholesteryl ester transfer protein (CETP) inhibitors and diabetes risk. Diabetes Metab 44, 508–513. 10.1016/j.diabet.2018.02.005.

