## Supplementary methods for "Insights into the metabolic consequences of type 2 diabetes"

### Quality control (QC)

#### UK Biobank (UKBB)

We removed individuals with high heterogeneity, sex mismatch, excess relatives, high relatedness and high missingness of the genotype based on the UKBB QC field (<https://biobank.ctsu.ox.ac.uk/crystal/crystal/docs/genotyping_qc.pdf>) . Further, for the purpose of our analysis, we used only White-British individuals (field 22006). All variants with a call rate below 98% and a p-value for Hardy-Weinberg equilibrium test below 10^-6^ were removed from the analysis. A total of 9,572,578 variants passed the QC.

The ukbnmr package was used to perform the metabolite QC on data at T0. After running the function remove_technical_variation() with the default parameters on the full set individuals with metabolite data (combining the first and second release), a total of 274,357 individuals were kept. In addition, we also removed individuals with a ‘Low Protein’ flag corresponding to individuals without enough material, leaving a total of 272,570 individuals, and we only kept individuals presenting less than 40% of missing data, leaving a total of 270,012 individuals. Finally, we restricted our analyses to the White British individuals defined from the field 22006 to work on a homogeneous population. The total number of individuals post-QC therefore corresponds to 227,607 individuals. No metabolites were removed from the analyses as all presented less than 1% of missing data (no imputation was performed on the missing levels).

#### Estonian Biobank (EstBB)

For the EstBB metabolite replication dataset, quality control and processing involved the removal of duplicated samples, removal of samples with more than 200 missing values, and donors who had withdrawn their consent. Prior analysis, inverse normal transformation was applied on the levels of each tested metabolite.

All EstBB participants have been genotyped using Illumina GSAv1.0, GSAv2.0, and GSAv2.0_EST arrays at the Core Genotyping Lab of the Institute of Genomics, University of Tartu. Individuals were excluded from the analysis if their call-rate was <95% or if their sex defined by heterozygosity of X chromosomes did not match their sex in the phenotype data. Pre-imputation quality control filters included call-rate <95%, Hardy–Weinberg equilibrium (HWE) P-value <1e^−4^, and minor allele frequency <1%. Pre-phasing was conducted using Eagle v2.3 software (https://www.ncbi.nlm.nih.gov/pmc/articles/PMC5096458/) and imputation was done using Beagle v.28Sep18.793 (https://www.sciencedirect.com/science/article/pii/S0002929718302428). The population-specific imputation panel of 2,297 whole genome sequencing samples was used (https://www.nature.com/articles/ejhg201751) as a reference. Related samples were removed from replication analyses. For that, IBD matrix was calculated by plink v1.90b6.21 (citation: https://academic.oup.com/gigascience/article/4/1/s13742-015-0047-8/2707533) with --genome setting. The custom R function was further used to remove related samples by using a pi-hat threshold of 0.15 while prioritizing T2D cases.

### Sensitivity analyses

BMI, lipid-lowering medication and metformin medication were sequentially considered as covariates in the interaction QTL and complications analyses as sensitivity analyses. BMI was extracted from the UKBB field 21001 was considered as a continuous variable. Medication information was extracted from the UKBB field 20003, and we selected the lipid-lowering medication from the ATC code groups C10A and C10B which was coded as a binary variable indicating whether the individuals were taking this type of medication or not. Information on metformin medication was retrieved from the UKBB field 20003 and was considered as a binary variable.
