## Supplementary material for "Insights into the metabolic consequences of type 2 diabetes": S1 Fig, S2 Fig, S3 Fig, S4 Fig

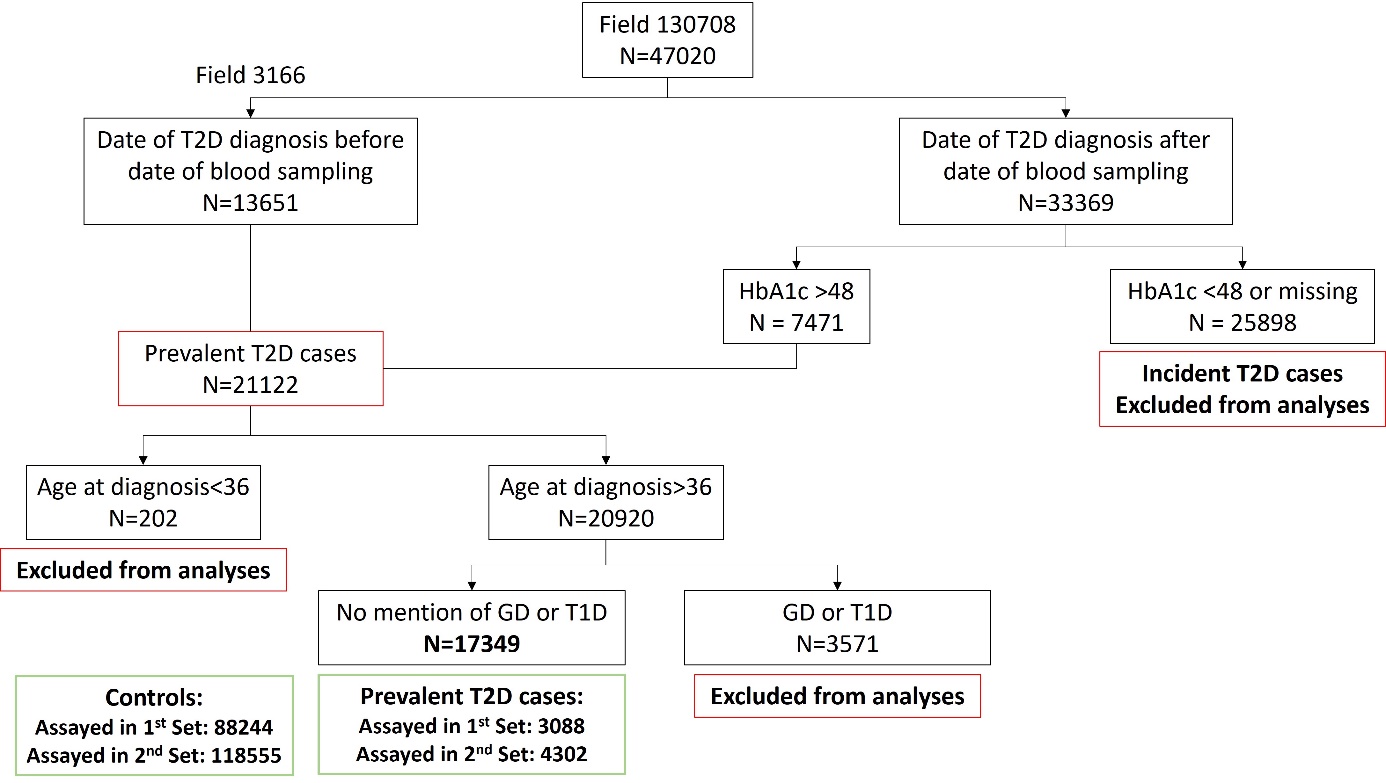

Figure S1: Workflow to define prevalent T2D cases at T0, when metabolite data have been measured.

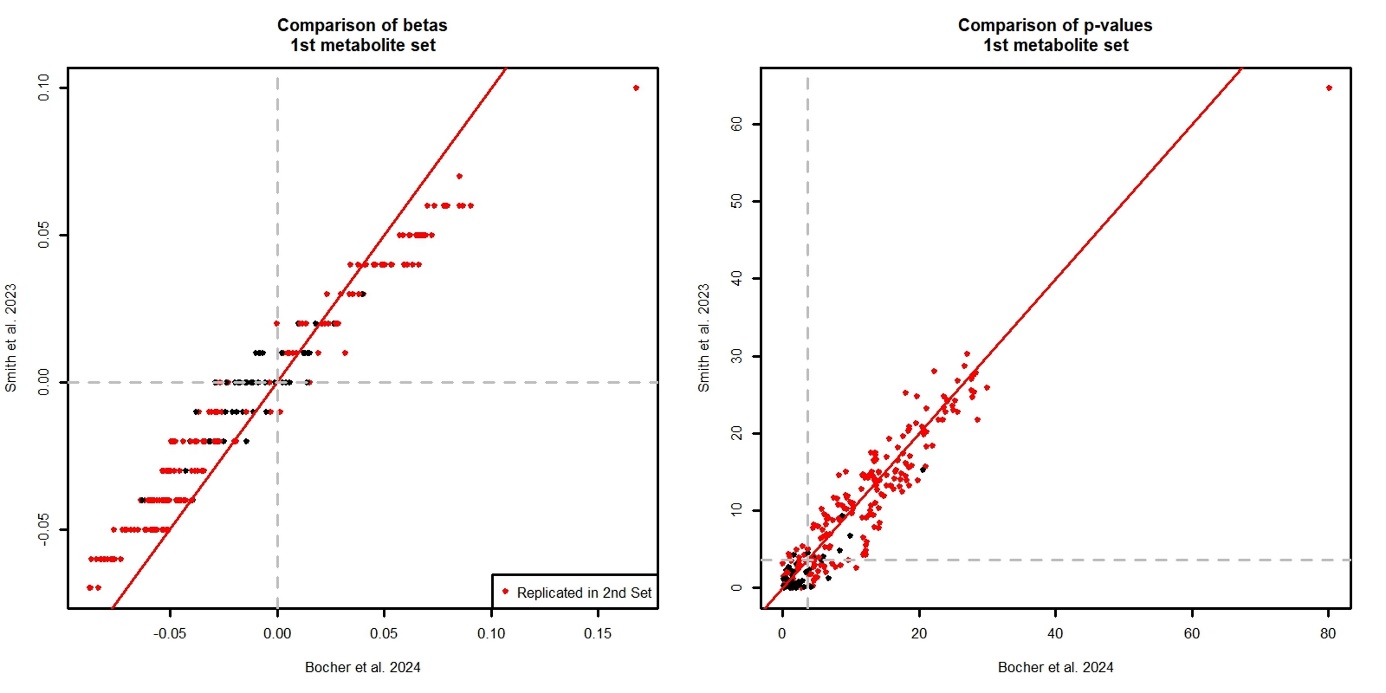

Figure S2: Comparison of betas and p-values with Smith et al. 2023 for the reverse MR on the first set of metabolites.

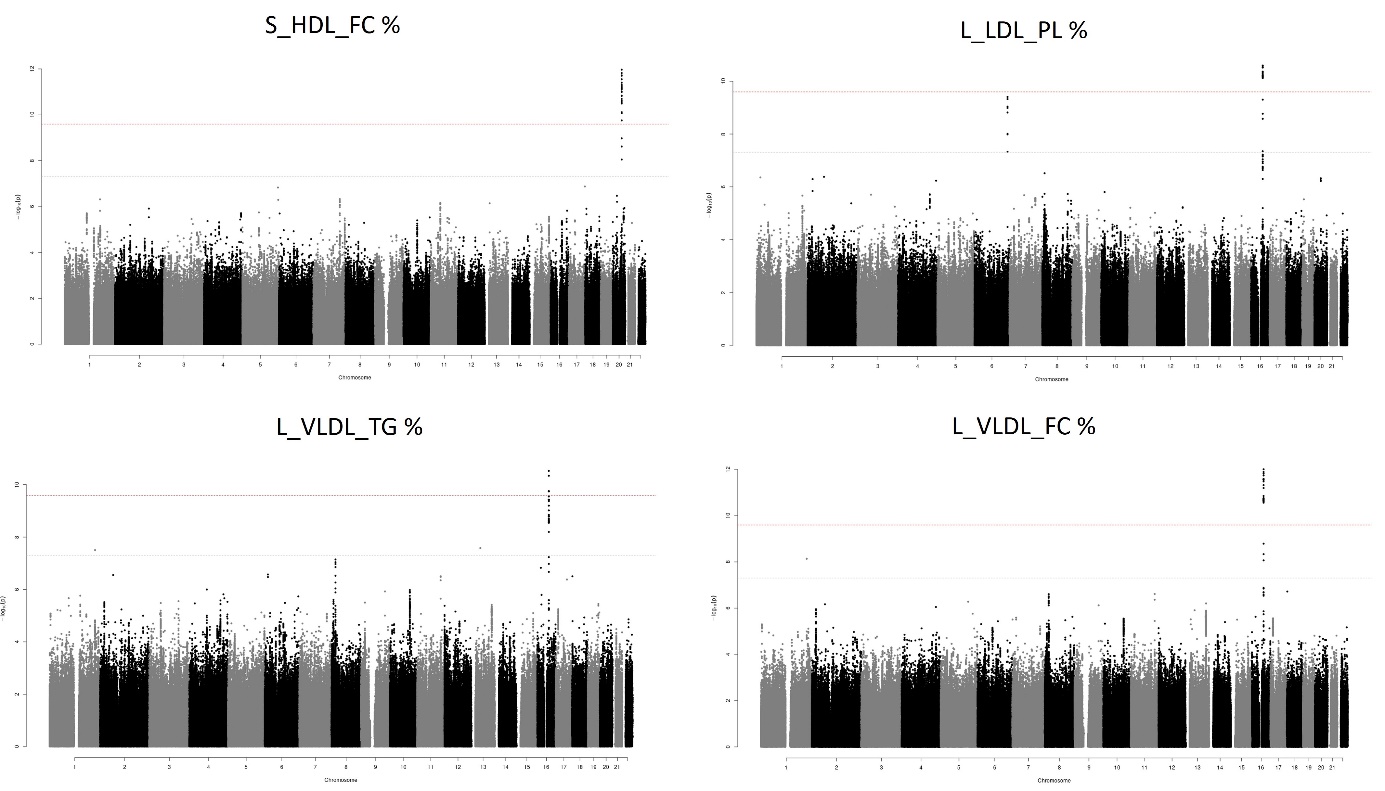

Figure S3: Manhattan plot of the interaction QTL analysis for the four metabolites significant and replicated in the EstBB.

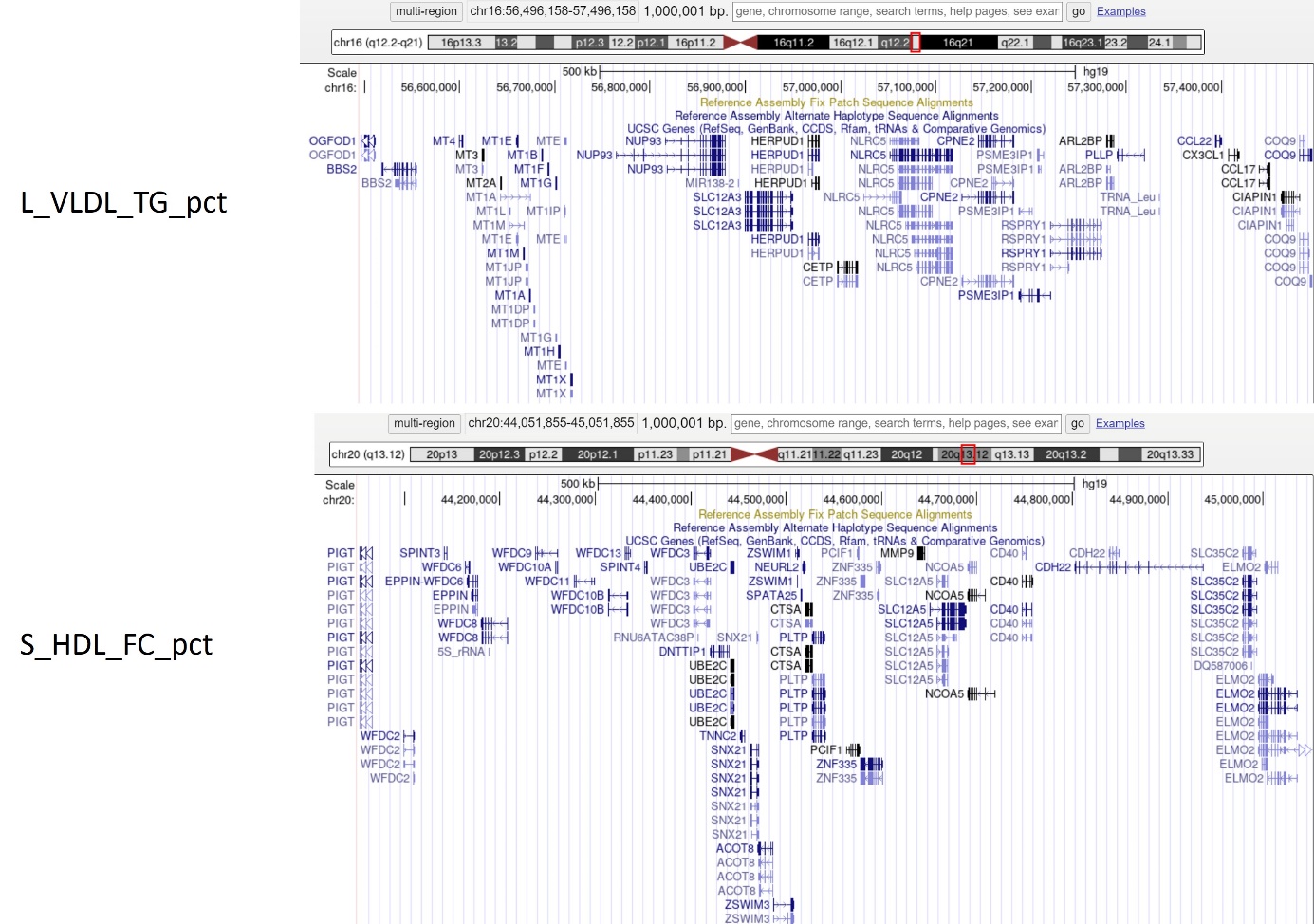

Figure S4: Genomic regions around the most significant variants from the interaction QTL analyses, obtained from https://genome.ucsc.edu/
